# Beyond Reorganization: Intrinsic cortical hierarchies constrain experience-dependent plasticity in sensory-deprived humans

**DOI:** 10.1101/2025.06.19.660510

**Authors:** Davide Orsenigo, Francesca Setti, Marco Pagani, Giovanni Petri, Marco Tamietto, Andrea Luppi, Emiliano Ricciardi

## Abstract

Innate cortical organisation and postnatal sensory experience interact dynamically to shape the functional architecture of the human brain. Using naturalistic stimulation, functional gradient analyses, and comparative approaches in congenitally blind, congenitally deaf, and typically developed individuals, we investigated how intrinsic hierarchical structures and sensory experiences influence cortical organisation. Our findings demonstrate that the principal functional gradients spanning from unimodal sensory to transmodal association cortices are consistently preserved across all groups, suggesting a robust genetically determined cortical scaffold. Nonetheless, congenital sensory deprivation selectively reshapes the geometry of modality-specific gradients, characterised by reduced functional differentiation within sensory-deprived cortical regions. These geometric contractions promote experience-driven plastic reorganization, enabling deprived sensory areas to establish enhanced functional connectivity with transmodal and non-deprived sensory cortices. Critically, this reorganisation aligns systematically with pre-existing cortical gradients, highlighting intrinsic hierarchical constraints that guide experience-dependent plasticity. Moreover, sensory-deprived regions exhibiting heightened connectivity actively engage in processing structured perceptual information from intact modalities, reflecting specific feature-driven cross-modal adaptations. Collectively, these results underscore a fundamental duality in cortical organisation: innate hierarchical principles impose constraints on cortical architecture, while sensory experience drives adaptive refinement, demonstrating the brain’s intrinsic capacity for flexible functional reconfiguration in response to sensory deprivation.

---

Understanding how sensory experience shapes brain development and functional organisation is central to neuroscience, bridging two long-standing theoretical perspectives. One view emphasises genetically encoded constraints as primary determinants of brain architecture [1, 2], while the other underscores the importance of experience-dependent plasticity in refining and adapting neural systems throughout development [3–5]. Although intrinsic structural and functional features of cortical organisation are evolutionary conserved across individuals and species, implying foundational principles of cortical architecture [6, 7], sensory experience significantly shapes the maturation and specialisation of neural circuits, particularly within modality-specific systems [8, 9]. This dynamic interplay between genetic predispositions and sensory experience becomes particularly evident in cases of early sensory deprivation, where substantial cortical reorganisation and connectivity provides unique insights into how experience modulates the development of large-scale neural networks [10–13].

Traditionally, investigation into brain organisation has primarily relied on discrete regional analyses and pairwise interactions under physiological or pathological conditions. However, recent advances underscore the importance of conceptualising the brain as a complex, economically optimised system operating across multiple spatial and temporal scales [14–18]. A significant limitation of region-based approaches lies in their inability to capture the flexible, context-dependent engagement of both primary sensory and higher-order transmodal cortices [19, 20]. In contrast, models describing cortical organisation along continuous gradients — from unimodal sensory regions to transmodal integrative areas — have garnered substantial empirical support, reflecting the brain’s inherent structural and functional heterogeneity [21–24].

Yet, the notion of cortical “hierarchy” itself encompasses diverse interpretations, including serial information flow, topological embedding, specialisation gradients, and developmental trajectories[25]. Converging evidence indicates that low-dimensional functional gradients, particularly a principal one extending from primary sensory/motor to higher-order transmodal cortices, effectively capture this multifaceted hierarchical architecture [26–35]. Notably, this principal gradient aligns with core developmental, evolutionary, and functional brain properties, such as prolonged postnatal maturation and progressive cortical specialization, supporting the emergence of higher-order cognition [27, 36].

While extensive research has characterised features of the principal gradient, how secondary gradients emerge and their modulation by sensory experience remain unclear. Empirical studies in congenitally blind or deaf individuals suggest that large-scale functional architectures can develop even without modality-specific sensory inputs, pointing to intrinsic scaffolds anchored in cortical geometry and structural connectivity [37–39]. Nevertheless, how experience-dependent mechanisms refine or reorganise these gradients, especially under sensory deprivation, remains unresolved [40].

To address these questions, we have analysed data from congenitally blind, congenitally deaf, and neurotypically developed participants presented with naturalistic stimuli (movie-watching of the 101 Dalmatians)[39]. This unique dataset enabled us to address two critical questions: (i) can cortical gradients emerge in the absence of modality-specific sensory input? and, (ii) How does sensory deprivation impact the spatial structure and functional specificity of these gradients?

We find that —while the fundamental hierarchical scaffold remains preserved regardless of sensory experience— deprivation leads to reduced specificity within the affected modality and fosters compensatory connectivity re-routing involving preserved unimodal and trans-modal regions. Collectively, these results highlight a dynamic interplay between innate cortical architecture and experience-dependent plasticity, suggesting that functional gradients are neither entirely hardwired, nor wholly experience-driven.

## RESULTS

### Gradient topography during processing of naturalistic auditory, visual and audiovisual processing in typically developed individuals

To characterise the macroscale functional architecture of the cortex under naturalistic stimulation, we computed pairwise functional connectivity (FC) matrices from regional BOLD time series, separately for each experimental group and condition. We then applied diffusion map embedding [41, 42] - a non-linear dimensionality reduction technique akin to principal component analysis (PCA;[16, 26, 43]) – to the group-level FC matrices to extract cortical gradients. We focused on the first three gradients, as they represent the primary hierarchical dimensions of functional organisation, spanning from unimodal sensory areas to transmodal association networks [26, 44].

Qualitative inspection of the extracted gradients revealed both shared structures and condition-specific topographies across experimental groups (Figure 2). Gradients derived from unimodal and multimodal naturalistic stimulation both reflect the canonical macroscale organisation of cortical hierarchy, and capture condition-dependent functional architectures. Consistent with prior findings [45], the embedding spaces under naturalistic stimulation showed enhanced differentiation along modality-specific axes, particularly within the visual and auditory systems (Figure 2A). The first gradient (G1) consistently reflects the canonical sensorimotor-to-transmodal cortical hierarchy across all conditions, mirroring intrinsic organisational principles also observed at rest [45]. However, in contrast to generalised embedding reported during rest, naturalistic stimulation elicited modality-specific refinements. During audiovisual and visual conditions, the second (G2) and third (G3) gradients were anchored in visual and auditory cortices, respectively, indicating the emergence of modality-specific granularity atop the preserved canonical structure. These findings suggest that naturalistic stimulation evokes distinct, context-sensitive cortical geometries, where dominant sensory modalities sculpt the expression of secondary gradients. Altogether, these results demonstrate that while the major organisational axes of the cortex are conserved across conditions, their relative prominence and spatial specificity are modulated by the sensory context. This indicates that functional gradients are dynamically shaped by naturalistic input.

**Figure 1.**
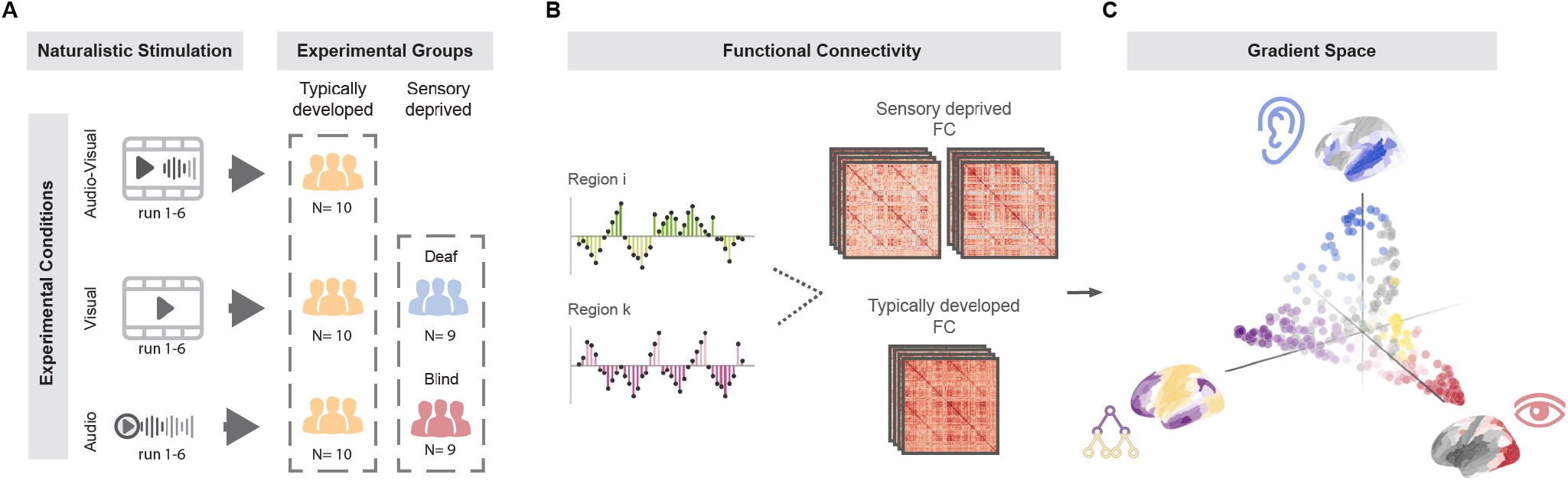
Experimental Paradigm and Data Analysis Pipeline. **A**, Participants viewed six runs of a naturalistic movie (101 Dalmatians) presented in three sensory formats: audiovisual, visual-only, and audio-only. The design included three groups of typically developed participants (N=10 per condition) and two groups of congenitally sensory-deprived individuals: blind (N=9) and deaf (N=8). **B**, Functional connectivity (FC) was computed from BOLD signals recorded via fMRI during movie viewing. FC matrices were estimated for each group and condition to examine pairwise regional interactions. **C**, Diffusion embedding was applied to the FC matrices to derive low-dimensional cortical gradients. These gradients were used to assess the global functional architecture and determine how congenital sensory deprivation affects its spatial geometry and hierarchical organization.

**Figure 2.**
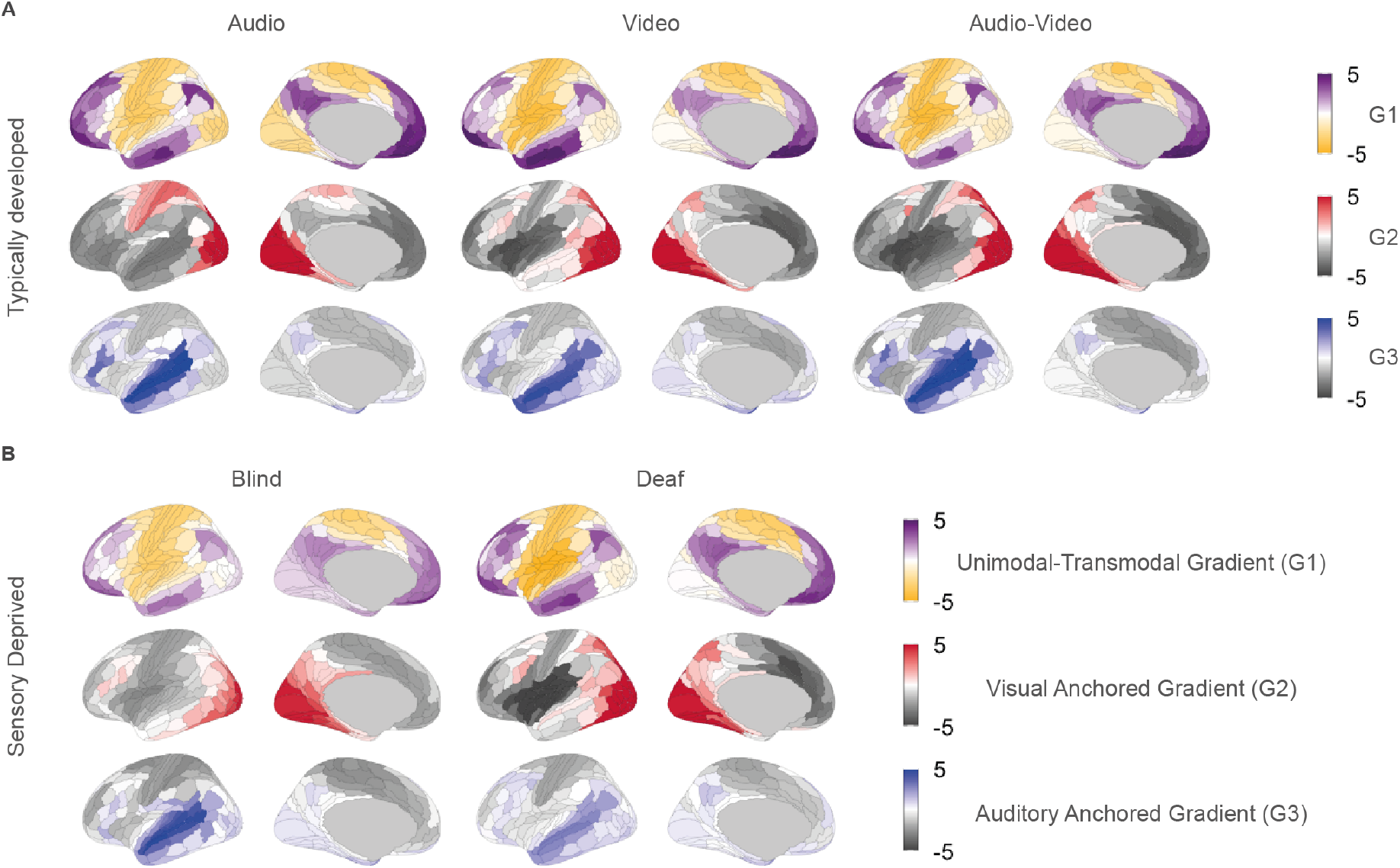
Topographical maps of functional gradients across groups and conditions. **A**, Surface representations of the first three cortical gradients are shown for typically developed participants under three naturalistic stimulation conditions: audio-only, visual-only, and audiovisual. Gradient 1 (G1) captures the principal unimodal-to-transmodal axis, while Gradient 2 (G2) and Gradient 3 (G3) are anchored in visual and auditory cortices, respectively. **B**, Corresponding gradient maps are shown for congenitally blind and congenitally deaf individuals, collapsed across sensory conditions. In both sensory-deprived groups, G1 remains consistent with the canonical cortical hierarchy, while G2 and G3 exhibit altered spatial differentiation, reflecting the absence of modality-specific experience..

### Sensory deprivation preserves gradient topography but reshapes functional cortical geometries

Building on our findings in neurotypically developed individuals, we next investigated how congenital sensory deprivation affects the expression and geometry of cortical gradients under naturalistic stimulation. If cortical gradients are intrinsically hardwired, they should be preserved regardless of sensory input. Alternatively, if postnatal experience is a constitutive prerequisite for shaping cortical organisation, we would expect marked differences in gradient structure under sensory deprivation. Our results reveal that the global hierarchical organisation of cortical gradients is largely preserved in both congenitally blind and deaf individuals. Specifically, the first three gradients consistently map onto a unimodal-to-transmodal axis (G1), a visually anchored axis (G2), and an auditory anchored axis (G3), respectively (Figure 2B). This preservation indicates that the macroscale scaffold of cortical functional geometry is established independently of modality-specific sensory experience.

However, we observed significant differences in the geometric properties of this low-dimensional embedding space. To do this, we leverage the range of a gradient, a simple, yet robust geometric measures, previously shown to be effective in capturing architectural changes across experimental conditions and species [46–49]. In particular, this metric reflects the spread and differentiation of connectivity profiles across cortical regions along one gradient. In this way, gradient range serves as a proxy for the degree of functional heterogeneity: greater ranges indicate more distinct, specialised regions, while reduced ranges signal functional homogenization. The unimodal-to-transmodal gradient range (G1) was preserved across groups compared to the audiovisual (AV) condition, suggesting that the principal gradient remains robust to sensory deprivation (two-tailed T-tests, blind vs. AV: T = -1.93, p = 0.07, FDR BH-corrected, Hedges’ g = -0.83; deaf vs. AV: T = 0.27, p = 0.79, FDR BH-corrected, Hedges’ g = -0.14; Figure 3B). In contrast, sensory deprivation led to significant contractions in the modality-specific gradients. In blind individuals, the range along the visual gradient (G2) was significantly reduced relative to both audiovisual and auditory conditions (blind vs. AV: T = –4.17, *p <* 0.001, FDR BH-corrected, Hedges’ g = –1.78; blind vs. A: T = –2.57, p = 0.02, FDR BH-corrected, Hedges’ g = –1.06; AV vs. A: T = 0.86, p = 0.39, FDR BH-corrected, Hedges’ g = 0.38). Similarly, a significant reduction in the range of the auditory gradient (G3) was observed exclusively in the deaf group, relative to the typically developed participants exposed to the video-only condition (deaf vs. AV: T = –5.6, *p <* 0.001, FDR BH-corrected, Hedges’ g = –2.31; deaf vs. V: T = –3.02, p = 0.01, FDR BH-corrected, Hedges’ g = –1.19; AV vs. V: T = 0.71, p = 0.48, FDR BH-corrected, Hedges’ g = 0.23). Altogether, these findings suggest that, while not mandatory for the emergence of macroscale gradient architecture, sensory experience plays a critical role in refining the functional differentiation of modality-specific systems. While the hierarchical organization of cortical gradients is preserved in congenital sensory deprivation, unimodal naturalistic stimulation reveals selective reductions in functional heterogeneity along the gradient corresponding to the deprived modality. This observation points to reduced differentiation in sensory-deprived areas, potentially rendering them more responsive to postnatal, experience-driven reorganization.

**Figure 3.**
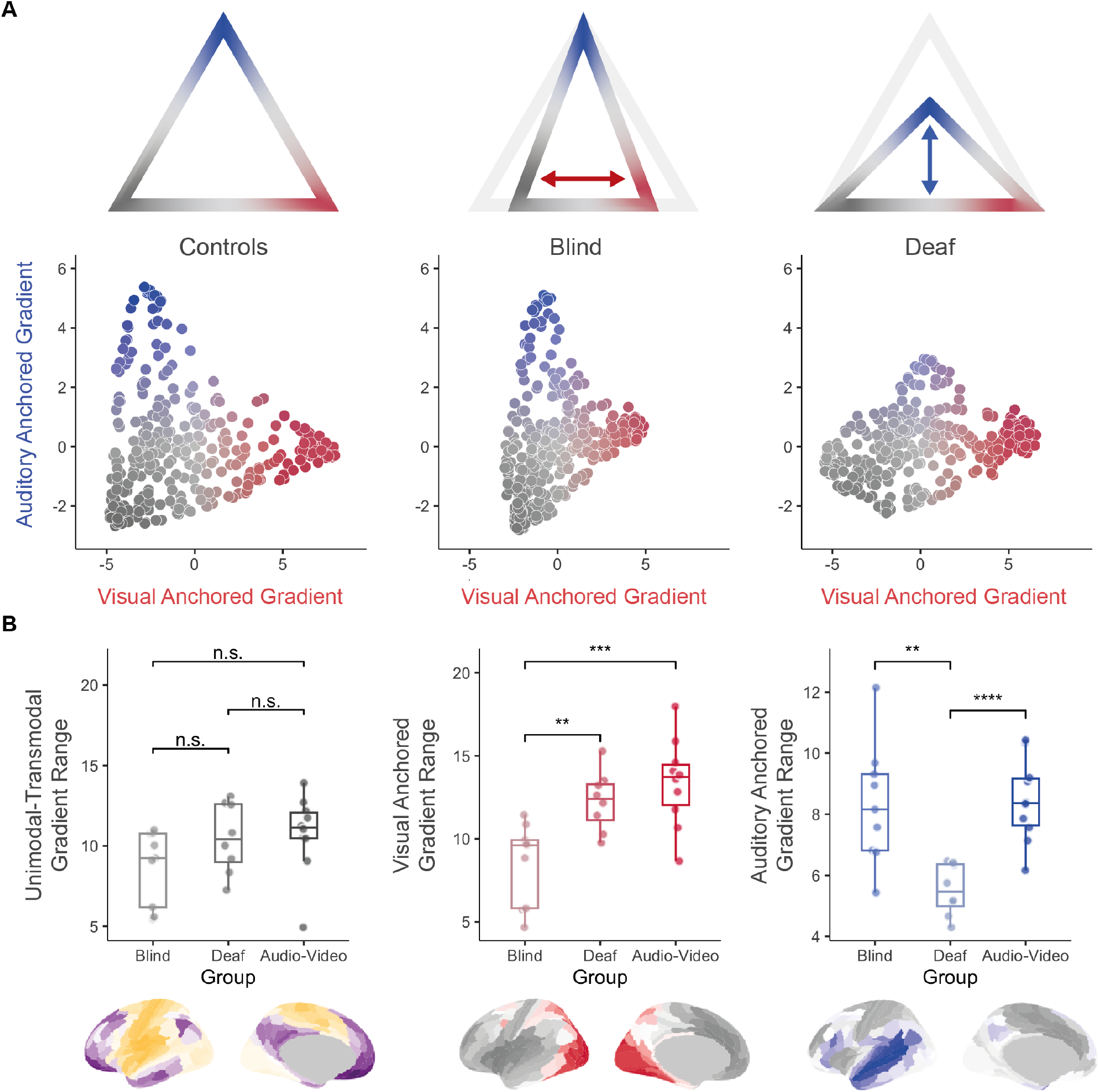
Functional geometry reconfiguration under sensory deprivation. **A**, Two-dimensional scatter plots illustrate the spatial embedding of cortical regions along the visual-anchored (G2) and auditory-anchored (G3) gradients in typically developed (audiovisual condition), congenitally blind, and congenitally deaf participants. The top panel schematically highlights the observed contraction patterns: visual gradient compression in blind individuals (horizontal arrow) and auditory gradient compression in deaf individuals (vertical arrow). **B**, Boxplots show gradient range values for the three groups along the first three functional gradients. No significant differences are observed for the unimodal-to-transmodal gradient (G1). In contrast, significant reductions in the visual gradient range (G2) are found in blind participants and in the auditory gradient range (G3) in deaf participants, compared to the typically developed group under audiovisual stimulation. Statistical comparisons are based on two-tailed t-tests with FDR BH correction (n.s. = not significant; ** *p <* 0.01; *** *p <* 0.001; **** *p <* 0.0001). Gradient maps below each plot depict the corresponding spatial topography.

### Functional Reorganization Driven by Sensory Gradients

To further explore the basis of gradient alterations under sensory deprivation, we analysed whole-brain functional connectivity patterns to assess how cortical interactions are reorganised. Treating functional connectivity (FC) matrices as undirected networks, we used the Network-Based Statistic (NBS; [50]) to identify significant differences in edge strength between groups (F=9, *p <* 0.05 corrected for multiple comparisons, FDR BH-corrected). For the blind group, FC matrices were compared to those of sighted individuals exposed to audiovisual and auditory-only movies. For the deaf group, comparisons were made against sighted participants exposed to audiovisual and visual-only conditions. This comparison scheme enabled us to isolate changes uniquely associated with congenital sensory deprivation rather than transient differences in unimodal stimulation. A composite null hypothesis test was applied to isolate differences shared across conditions [51](Figure 4A). Results showed that reorganisation in both groups primarily involved enhanced connectivity between unimodal and transmodal areas, rather than diffuse connectivity changes. (i.e., greater Pearson correlation between their BOLD timecourses) (see Figure 4A). These effects were consistently more prominent in blind individuals (1% of statistically significant connections altered) than in deaf individuals (0.001% altered). This approach revealed that congenitally blind individuals exhibit increased connectivity between deprived extrastriate visual areas (e.g., MT+ complex and dorsal stream cortices) and the dorsolateral and inferior frontal cortices (frontoparietal and DMN regions), alongside decrease in intra-visual network connectivity. Congenitally deaf participants showed increased connectivity between extrastriate visual areas (including the MT+ complex and neighbouring regions) and auditory association cortices (e.g., STS and higher-order auditory areas A4 and A5). Recent work indicates that intrinsic activity and connectivity mature along a hierarchical sensorimotor–association axis, with unimodal and transmodal regions developing along distinct trajectories. In turn, this suggests that such an intrinsic hierarchy may shape processes of experience-dependent functional reorganization [52, 53]. Following this hypothesis and having first shown that individuals with congenital blindness and deafness exhibit a contraction of the gradients corresponding to the deprived sensory modality, we now investigate whether the observed experience-dependent reorganization in functional connectivity is somehow ‘constrained’ and unfolds along the intrinsic sensory gradients themselves.

We correlated statistical maps of FC differences with the corresponding sensory gradients computed in the typically developed audiovisual group. Thus, checking the correlation between the average involvement of brain areas in the interactions elicited by the sensory deprivation and their position along the corresponding sensory gradient (i.e., the second one for the blind population, the third one for the deaf population) computed on the typically developed audio-video population (Figure 4C). Strikingly, for both blind and deaf individuals, significant positive correlations were observed between the spatial pattern of reorganisation and the corresponding sensory gradient, even after correction for spatial autocorrelation (blind - visual gradient; Spearman rho = 0.42, *p <* 0.001, *p*_*null*_ *<* 0.001; deaf auditory gradient; Spearman rho = 0.36, *p <* 0.001, *p*_*null*_ = 0.003). This suggests that compensatory connectivity changes in congenitally deprived unimodal cortices unfold along pre-existing functional gradients. Of note, this observation favors that intrinsic functional architecture preserves not only large-scale portions of cortical organization, but also ‘chaperons’ experience-dependent reorganization. To understand the functional relevance of these reorganised networks, we next examined how BOLD activity in regions showing increased FC relates to perceptual content. We correlated BOLD time courses with annotated features of the movie stimulus, ranging from low-level visual (e.g., static Gabor-like filters and motion energy from spatiotemporal integration) and auditory features (e.g., spectral profiles and amplitude envelopes), to mid-level speech-related attributes (e.g., word and letter counts, dialogue presence) and higher-level semantic content (e.g., categories of natural vs. artificial elements in auditory and visual streams, and word embeddings derived from subtitles) [39]. We then tested for group differences in these correlations to determine whether reorganised regions process distinct content. Results revealed significant group differences in correlations between distinct movie features and activity within regions previously identified as exhibiting increased functional connectivity. Specifically, congenitally deaf participants showed stronger correlations between auditory association cortices and visual movie features compared to the AV control group, while blind individuals showed stronger correlations between deprived visual areas and mid-level speech features compared to AV controls (two-tailed T-tests, FDR BH-corrected see Supplementary Table SX; significant effects displayed in Figure S12). Thus, we confirm that auditory and speech-related features are redirected to deprived visual cortices in blind individuals, while visual features are rerouted to deprived auditory cortices in deaf individuals, reflecting structured, modality-specific cross-modal reorganization. Altogether, these findings demonstrate that functional reorganization after congenital sensory deprivation adheres to intrinsic cortical hierarchies and supports structured cross-modal processing. Deprived sensory areas do not become functionally dormant; rather, they are recruited for computationally relevant roles through experience-dependent reconfiguration constrained by the brain’s inherent architecture.

**Figure 4.**
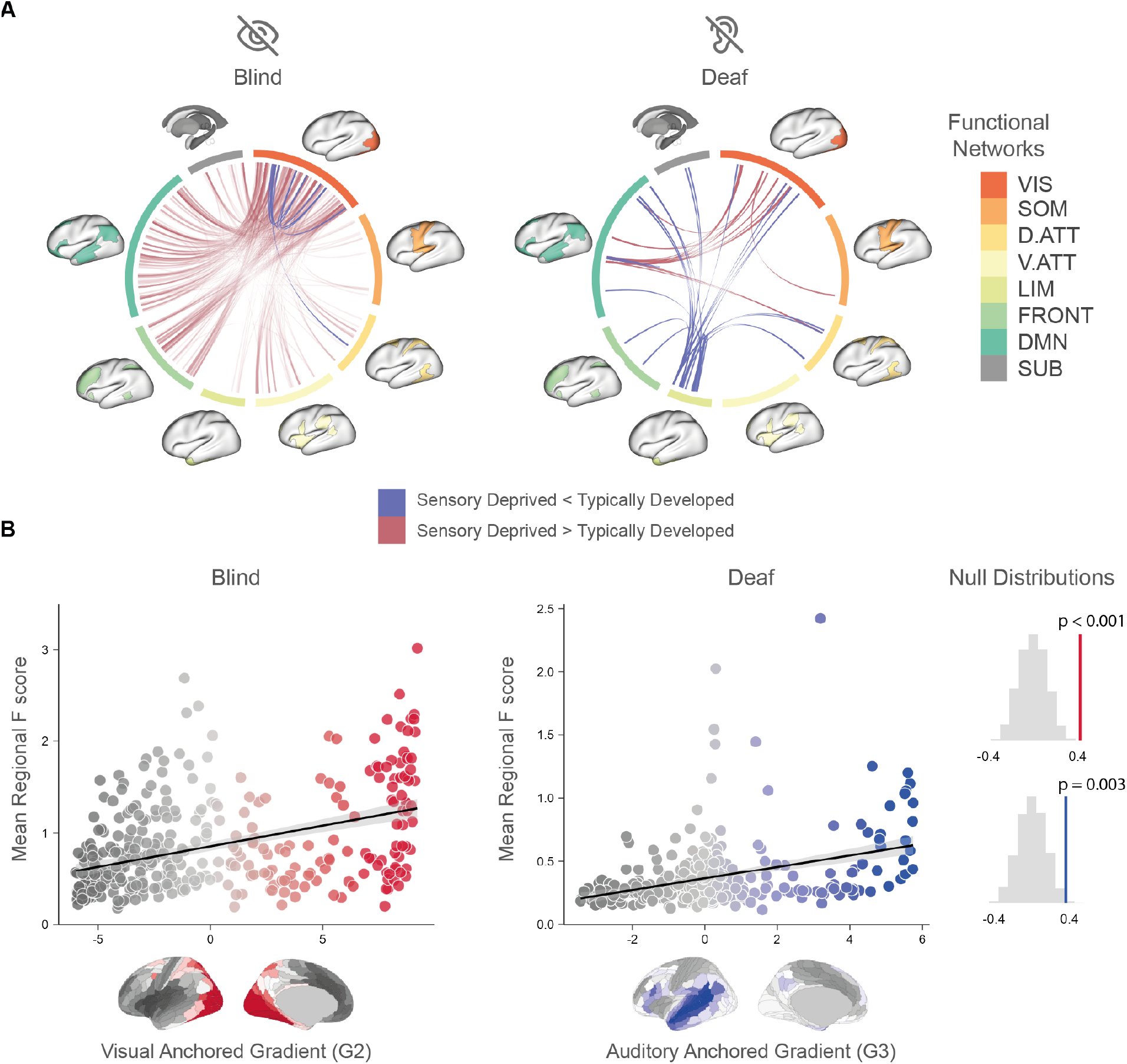
Functional reconfiguration follows intrinsic cortical hierarchies. **A**, Group-level differences in whole-brain functional connectivity between congenitally blind or deaf individuals and matched typically developed controls were identified using the Network-Based Statistic (NBS; *F* ≥ 9, *p <* 0.05). Chord diagrams illustrate significantly increased (blue) and decreased (red) connections in each sensory-deprived group. Cortical nodes are grouped by canonical functional networks [54]: VIS = visual, SOM = somatomotor, D.ATT = dorsal attention, V.ATT = ventral attention, LIM = limbic, FRONT = frontoparietal, DMN = default mode, SUB = subcortical. **B**, Mean regional F scores from the NBS analysis were correlated with the spatial layout of the visual-anchored gradient (G2) in the blind group and the auditory-anchored gradient (G3) in the deaf group. Significant positive correlations were observed in both groups, even after correcting for spatial autocorrelation using null distributions derived from 10,000 spatially-constrained surrogate maps (*p <* 0.001 and p = 0.003, respectively; right panel)(Markello and Misic [55], Burt *et al*. [56]; see Methods). This analysis demonstrates that functional reorganization in sensory deprivation unfolds along pre-existing cortical gradients.

**Figure 5.**
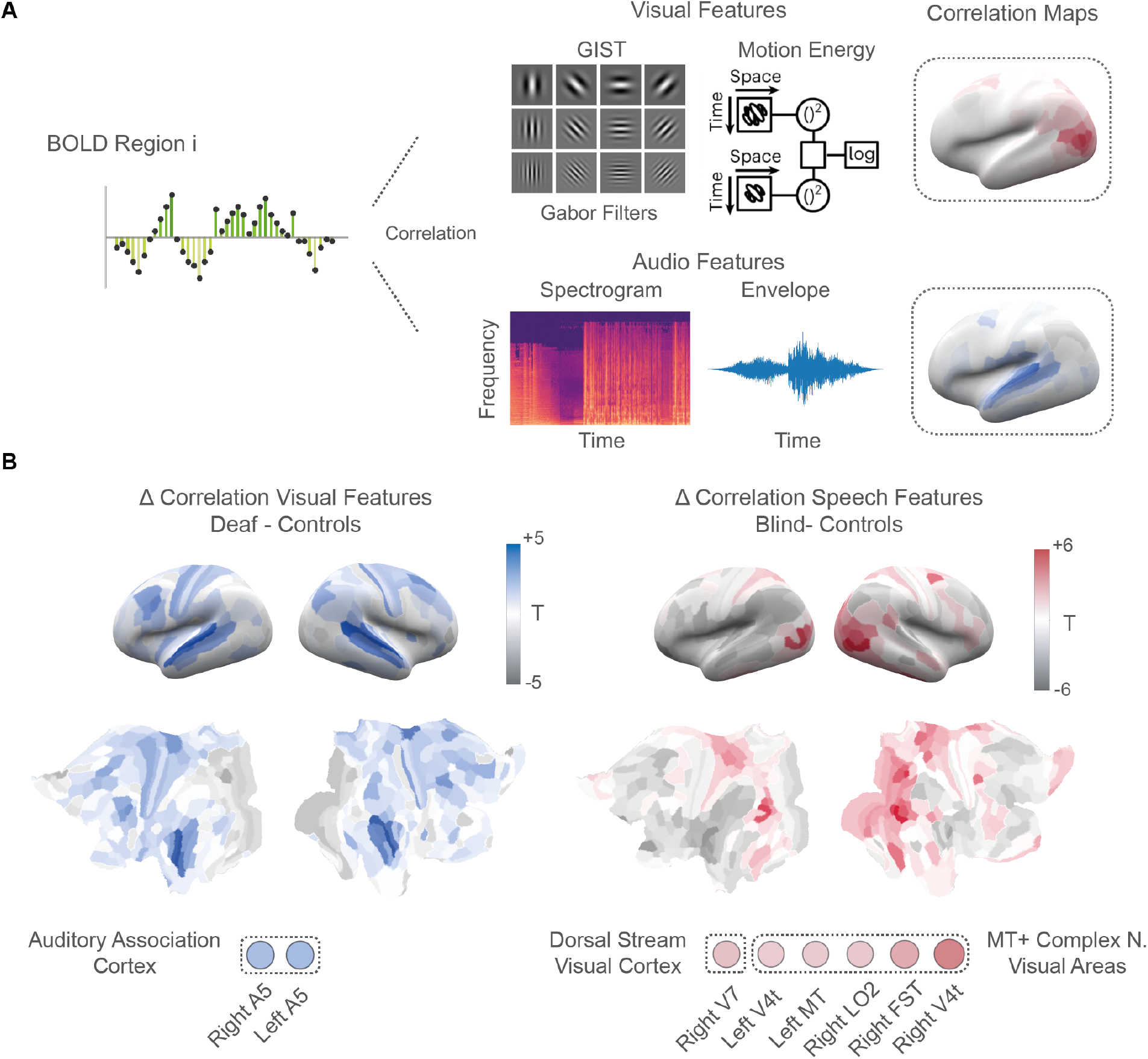
Feature-specific recruitment of deprived sensory cortices. **A**, Schematic of the analytical pipeline: BOLD time series from each cortical region were correlated with time courses of annotated stimulus features, including low-level visual (e.g., Gabor filters, motion energy), low- and mid-level auditory features (e.g., spectrogram, envelope), and speech-related semantic features. Correlation maps were generated per subject and compared across groups **B**, Surface renderings and inflated ROI maps show results from region-wise two-tailed t-tests contrasting sensory-deprived and typically developed individuals. Left: Deaf participants exhibit significantly stronger correlations between visual stimulus features and auditory association cortices (blue, *p <* 0.05 FDR BH-corrected). Right: Blind participants show significantly stronger correlations between speech-related features and visual cortices, including MT+ and dorsal stream regions (red, *p <* 0.05 FDR BH-corrected). Insets highlight significant clusters and annotated ROIs.

### Robustness and Sensitivity

To assess robustness to the choice of free parameters—specifically the sparsity and diffusion parameters used to compute the functional gradients—the main analysis was replicated across *α* values of 0.3, 0.7, and 0.9 and sparsity levels of 70% and 80%. The analysis was then repeated using an alternative parcellation, namely the 400-region functional atlas of Schaefer *et al*. [57], which yielded coherent results preserving the number of relevant interactions. We further evaluated the sensitivity of the NBS findings to a stricter F-threshold (*F* = 12), confirming the stability of our results. Finally, the stability of the feature-correlation analysis was assessed by replacing word counts with a binary indicator of dialogue presence in each TR.

## DISCUSSION

How do sensory experience and innate organisational principles jointly shape the macroscale functional architecture of the cortex? Here, leveraging naturalistic stimulation and functional gradient analyses, we compare three distinct sensory conditions—audiovisual, visual-only, and auditory-only— in congenitally blind, congenitally deaf, and neurotypically developed individuals to examine whether sensory deprivation affects hierarchies and functional gradients.

We demonstrate that congenital sensory deprivation does not disrupt the overall hierarchical organization of cortical functional gradients. However, sensory-deprived cortices exhibit reduced functional differentiation, in particular along the gradient dimension associated with the deprived modality. This reduced differentiation appears to facilitate postnatal, experience-dependent modelling of deprived cortical areas. Notably, such reorganization unfolds along the intrinsic sensory gradients themselves, with perceptual information from spared unimodal or transmodal regions being systematically rerouted to sensory-deprived areas. Altogether, these results underscore that the development of cortical architecture emerges from a dynamic interplay between intrinsic, genetically driven organizational frameworks and extrinsic, experience-dependent shaping forces.

Brain macroscale functional organization appears both to be anchored by innate hierarchies that constrain plasticity, yet sufficiently flexible to accommodate profound changes in sensory experience from the earliest stages of development. Extending recent observations by Samara *et al*. [45], our data indicate that during naturalistic processing, the primary gradient retains its canonical resting-state configuration, spanning from unimodal sensory to transmodal regions, independent of sensory experience.

This robust preservation aligns with evolutionary and developmental evidence that suggest an intrinsic organisational scaffold underpinning large-scale functional architecture [27]. Indeed, prior studies support the notion that the principal gradient may reflect fundamental organising principles anchored by intrinsic factors, such as gene expression patterns and cortical microstructure, transcending specific sensory modalities [12, 39, 58–63]. In contrast, we observed selective alterations in the geometric properties (i.e., contractions in the gradient range) of the secondary gradients corresponding to deprived sensory modalities. Gradient range has previously been established as a sensitive biomarker of functional heterogeneity across the cortical surface, with broader ranges denoting greater differentiation, and narrower ranges reflecting more homogeneous functional architectures [46–49].

In the context of congenital deprivation, reduced range in visual or auditory gradients likely reflects a disruption of this maturational refinement process, whereby modality-specific input is typically required to sculpt the fine-grained functional topography of unimodal areas. Importantly, this reduction is not global but modality-specific, selectively affecting the gradient dimension anchored in the deprived sensory cortex. The modality-specific nature of this gradient contraction highlights their sensitivity as indicators of experience-driven differentiation.

Our findings support a dual-mechanism model wherein intrinsic, genetically guided structural constraints establish a stable global hierarchical scaffold, while sensory experience fine-tunes local cortical specialisation [16, 25, 40, 48]. This dual influence of structural anchoring and sensory-driven refinement closely aligns with recent theoretical models suggesting that functional specialization is not solely dictated by sensory input, but also by the computational goals of each cortical area, its embeddedness in pre-existing anatomical and functional networks and experience-dependent adaptation [39, 64– 68].

Our gradient-based findings are complemented by finer-grained connectivity analyses revealing modality-specific reorganization patterns. Previous evidence showed modified functional connectivity between deprived unimodal cortices and non-deprived unimodal or transmodal areas during sensory processing (e.g., Amaral *et al*. [59], Anurova *et al*. [69], Bauer *et al*. [70], Amaral *et al*. [71]). Consistently, in congenitally blind individuals, we observe increased connectivity between extrastriate and key hubs of the frontoparietal and default mode networks.

This increased visual-to-frontal integration occurs alongside a reduction in intra-network connectivity within the visual system itself, suggesting a reorganization of functional resources toward higher-order transmodal regions. Additionally, the emergent connectivity between the deprived visual cortex and the default mode network in blind individuals mirrors the functional reorganization observed in sensorimotor cortex after limb loss, suggesting that, in the absence of primary input, cortical regions may be reallocated to internally driven functions through integration with transmodal networks [72].

A complementary pattern emerges in congenitally deaf individuals, who exhibited strengthened connectivity between the visual network and auditory association cortices, notably the superior temporal sulcus and high-level auditory regions. Therefore, our findings confirm that deprived unimodal cortices develop reorganized, structured interactions with functionally and topographically distant regions. By testing the relationship between the spatial distribution of functional changes and the underlying sensory hierarchy, we also showed that this functional reorganization follows the pre-existing sensory hierarchies of the brain. In other words, deprived cortical regions tend to establish new connections with areas that already lie along specific functional gradients, reflecting the typical sensory organization.

This implies that the brain’s intrinsic architecture not only remains intact in the absence of sensory input, but also actively guides plastic reorganization along inherent large-scale structure. Consequently, the altered connectivity patterns in blind and deaf people provide compelling support for the idea that functional reorganization unfolds along meaningful computational and anatomical trajectories, and highlight how intrinsic geometry and long-range pathways enable the deprived sensory cortex not only to remain functionally relevant but to contribute adaptively to non-canonical information processing through experience-dependent reconfiguration.

Importantly, these reorganized interactions appear to reflect specific patterns of feature-oriented integration of perceptual information on low/mid-level features. Our data also reveal a distinctive pattern of functional recruitment that differs fundamentally between blindness and deafness, pointing to important differences in the reorganizational potential of the occipital and temporal cortices. The occipital cortex’s extensive intrinsic and extrinsic connections, early maturation, and central hierarchical positioning confer exceptional versatility and computational flexibility, facilitating significant functional repurposing in blind individuals [12, 68, 73, 74]. In contrast, the auditory cortex demonstrates relatively constrained reorganizational potential, relying primarily on strengthening pre-existing heteromodal connections modulated by top-down influences from transmodal regions [75].

In the light of our results, we propose that intrinsic architectural functional geometry governs spatially extended cross-modal plasticity, thus providing a unified model explaining both stability and flexibility in sensory-deprived brains. Explicitly testing the evolution of chronic readjustment, longitudinal studies across different developmental stages (including late-onset sensory deprivation) will be invaluable to determine whether functional changes occur within a critical period or persist beyond it. Recent evidence from congenitally sensory-deprived individuals shows that meaningful, content-specific representations can emerge independently of a postnatal experience, pointing to an organizational blueprint or proto-architecture that predates and scaffolds experience-dependent specialization (e.g., Ricciardi and Pietrini [12], Setti *et al*. [39], Arcaro and Livingstone [64], Ricciardi *et al*. [68]). Notably, supramodal processing — where cortical regions encode conceptual content regardless of sensory modality — has been extensively demonstrated and supports the idea of modality-independent representational formats across sensorimotor experiences and higher cognition representations [12, 68].

Moreover, this perspective aligns with evidence that even primary sensory areas, traditionally viewed as unimodal, are embedded in a richly interconnected architecture that supports early multisensory integration. Unimodal cortices may exhibit the capacity to integrate cross-modal input that becomes functionally dominant when the canonical sensory input is absent, as often seen in sensory deprivation [76, 77]. In addition, our data could also be integrated into the framework arguing that the brain’s functional architecture is structured to encode and respond to features of sensory experience that transcend any single modality.

In this view, the dialogue across modalities, rather than a mere compensatory adaptation, reflects an intrinsic, supramodal infrastructure that allows the brain to flexibly route, compare, and integrate inputs from different sensory systems to detect and respond to environmentally relevant changes (e.g., Novembre and Iannetti [78]).

In summary, our study highlights that the macroscale functional architecture of the cortex emerges from a dynamic interplay between genetically guided intrinsic constraints and extrinsic sensory experience. Functional gradients provide a valuable framework to understand this dual influence, illustrating how innate structural scaffolds impose consistent organisational principles while remaining sufficiently plastic to accommodate adaptive changes. Our findings demonstrate that even under conditions of extreme sensory input loss, cortical reorganisation remains anchored within existing hierarchical structures, underscoring the presence of a stable cortical blueprint. Brain macroscale functional organisation appears both to be anchored by innate hierarchies that constrain plasticity, yet sufficiently flexible to accommodate profound changes in sensory experience from the earliest stages of development.

## MATERIALS AND METHODS

### Ethical statement

We took advantage of a previously acquired dataset, and full details are provided in the original publication [39]. Each volunteer was instructed about the nature of the research and gave written informed consent for participation, in accordance with the guidelines of the institutional board of the Turin University, Brain Imaging Centre. The study was approved by the Ethical Committee of the University of Turin (protocol number 195874, May 29th 2019) and conforms to the Declaration of Helsinki. Data used in this analysis were drawn from the study by Setti *et al*. [39], meaning the participant sample, stimulation, data acquisition and preprocessing is identical to the one described in the original work, as outlined in the following paragraphs.

### Participants

Fifty participants took part in the study. As already documented in Setti *et al*. [39], thirty normally-developed (TD) individuals and twenty sensory-deprived (SD) subjects (born without visual or auditory experience) were enrolled. The TD group was split into three equal samples, each exposed to one version of the same feature film: (1) full audiovisual (AV), *N* = 10 (age 35 ± 13yr; 8 females), (2) auditory only (A), *N* = 10 (age 39 ± 17yr; 7 females), (3) visual only (V), *N* = 10 (age 37 ± 15yr; 5 females). The SD cohort comprised congenitally blind participants (*N* = 11, mean age 46 ± 14yr; 3 females) who viewed the A version, and congenitally deaf participants (*N* = 9, mean age 24 ± 4yr; 5 females) who viewed the V version. Two blind subjects were excluded from the fMRI analyses due to excessive head motion (final SD sample: *N* = 9, mean age 44 ± 14yr; 3 females). All participants were right-handed according to the Edinburgh Handedness Inventory, native Italian speakers, and had no history of neurological or psychiatric disorders. The deaf group was proficient in Italian Sign Language and did not use hearing aids; the TD group reported normal or corrected-to-normal vision, intact hearing, and no knowledge of sign language. Further details for the SD participants are provided in Supplementary Materials of Setti *et al*. [39].

### Naturalistic stimulus

Following Setti *et al*. [39], naturalistic stimulation was provided via three versions—visual (V), auditory (A) and audiovisual (AV)—of the live-action film *101 Dalmatians* (Herek, Great Oaks Entertainment & Walt Disney, 1996). To facilitate compliance during unimodal presentation, a linear-plot narrative was selected. The original film was edited to fit a single scanning session by omitting non-essential scenes and splicing remaining footage to maintain narrative continuity. The final cut spans approximately 54 minutes and is divided into six runs of ∼ 8 minutes each, each preceded and followed by a 6 s fade-in and fade-out.

For the A condition, an Italian audio description was professionally recorded and overlaid on the film’s original soundtrack to convey omitted visual content. The script was adapted to bridge edited gaps and to describe essential visual elements not captured by dialogue or music. Recording was performed in a noise-isolated studio using a Neumann U87 ai microphone, Universal Audio LA 610 mk2 preamplifier, Apogee Rosetta converter, and Logic Pro 10.4. The voice track was then mixed with the original audio, and fade effects were applied to smooth transitions between runs; music and narration were subsequently remixed for level balance.

The complete soundscape (dialogue, narration, environmental sounds) was transcribed into styleand colorcoded subtitles to distinguish speaking voices and aid comprehension. Line breaks were adjusted (oneor twoline formats) to avoid interference with reading or visual processing. Video editing was carried out in iMovie (v10.1.10), and subtitles were authored using Aegisub (v3.2.2, http://www.aegisub.org/). In the V and AV conditions, a small red fixation cross was superimposed at screen center, with subtitles displayed at the bottom.

### fMRI experimental design

Prior to scanning, all participants rated their familiarity with the film’s plot on a 5-point Likert scale (1 = not at all, 5 = very well). During fMRI acquisition, each experimental subgroup viewed one of the edited versions of the film (V, A or AV) and were instructed simply to enjoy the presentation. Structural and functional MRI data were collected in a single session. Upon completion of imaging, engagement and compliance were evaluated using a custom two-alternative forced-choice questionnaire probing key narrative elements.

### Stimulation setup

Audio and visual stimuli were presented via MR-compatible LCD goggles and headphones (VisualStim Resonance Technology; video resolution 800×600 at 60Hz; visual field 30°×22°; 5” display; audio attenuation 30dB; frequency response 40Hz–40kHz). Both devices were supplied to all participants regardless of experimental condition. Stimulus delivery was managed using Presentation 16.5 (Neurobehavioral Systems; http://www.neurobs.com).

### fMRI data acquisition and preprocessing

Functional MRI data were acquired on a Philips 3T Ingenia scanner equipped with a 32-channel head coil. Gradient-echo EPI sequences were used with the following parameters: TR = 2000 ms; TE = 30 ms; FA = 75°; FOV = 240 mm; matrix = 80×80; slice thickness = 3 mm; voxel size = 3×3×3 mm; 38 axial slices acquired in ascending order; 1614 volumes were collected across the six movie runs, with an additional 256 volumes for the scrambled control run. High-resolution T1-weighted anatomical images were obtained using an MPRAGE sequence (TR=7ms; TE=3.2ms; FA=9°; FOV=224mm; matrix=224×224; slice thickness=1mm; voxel size=1×1×1mm; 156 sagittal slices). Data acquisition and analysis were conducted with full knowledge of experimental conditions.

Preprocessing followed standard AFNI (v17.1.12) workflows. First, spike artifacts were removed (3dDespike), and all volumes within each run were slice-time corrected (3dTshift) and realigned to the first volume of the first run (3dvolreg). Spatial smoothing was applied using a 6mm FWHM Gaussian kernel (3dBlurToFWHM), followed by percent signal normalization. Each run was detrended using a Savitzky–Golay filter (polynomial order=3, frame length=200 timepoints) in MATLAB R2019b to remove low-frequency drifts and outliers. The runs were then concatenated, and nuisance regressors—including motion parameters and spike regressors for framewise displacements ¿0.3mm—were removed via multiple regression (3dDeconvolve). Finally, individual participant time series were nonlinearly warped to MNI-192 standard space using 3dQWarp.

### Movie annotations

In the previous work by Setti *et al*. [39], the authors derived a set of movie-related features by means of computational modeling. Among these, low-level features were characterized both for the auditory and visual streams, thus capturing spectral (frequency) and sound envelope properties on one hand and static Gabor-like filters (GIST) and motion energy information on the other. Additionally, high-level features were defined based on manually tagged natural and artificial categories as well as word embeddings from subtitles. For the latter, two alternative embeddings were developed: one from individual sentences, utilizing the pretrained English-based GPT-3 model to fully capture semantic compositionality, and one using single-word embeddings generated with the Word2Vec algorithm trained on an Italian corpus. This produced two high-level semantic models, one combining categorical data with GPT-3 embeddings and the other with Word2Vec embeddings. Features related to the film’s editing were also annotated (such as scene transitions, cuts, dialogues, music, and audio descriptions). To remove shared variance across models, each stimulus model was orthogonalized with respect to movie editing descriptors. The editing features (e.g., cuts, transitions, dialogues, and music) were shown to influence both low-level descriptors (e.g., shifts in visual and auditory properties at scene transitions) and high-level semantic descriptors (e.g., spoken and written dialogues), potentially masking the finer computational features and inflating the explained variance. The methodological approach and results are detailed in Supplementary Information of the original work. Based on the movie script, two new speech-specific annotations were created. The script was segmented according to the repetition time (TR) of the functional images, with each time window annotated to indicate the binary presence or absence of dialogue. Additionally, a letter count within each segment was included to model the presence of human speech at a low level of complexity.

### Brain Parcellations

Cortical segmentation was performed using the 360-region multimodal parcellation of Glasser *et al*. [7], which provides functional and anatomical cortical mapping. Within each ROI, denoised BOLD-signal time series were averaged across all voxels to yield a single representative time course. These ROI-wise time courses were then extracted for subsequent analyses. One participant in the deaf cohort was excluded because the R TGv region lay almost entirely outside the acquired volume.

### Analytical Pipeline

#### Functional gradients from diffusion map embedding

Cortical functional gradients were calculated using the BrainSpace toolbox (https://github.com/MICAMNI/BrainSpace) in Python, applying default settings for kernel sparsity estimation (0.9), similarity kernel (cosine similarity), and anisotropic diffusion parameter (*α*=0.5). We calculated the functional connectivity matrix (FC) computing for each ROI s of the HCP Atlas [7] the Pearson correlation with all the other cortical areas. The obtained matrix was then row-wise z-transformed and thresholded to obtain a 10% density, thus retaining only the strongest interactions and removing possible spurious effects. Given the resultant matrix, a normalized cosine angle affinity matrix is computed, indicating the similarity in connectivity patterns for each pair of ROIs. Finally, the obtained affinity matrix is embedded in a low-dimensional space by means of diffusion map embedding. This approach has the advantage, compared to other approaches like PCA, of being non-linear and thus it allows to account for more complex patterns. The algorithm is governed by parameters *α* and t, where *α* determines the influence of the density of sampling points on the manifold (*α* = 0 indicates maximal influence, while *α* = 1 indicates none) and it adjusts the scale of eigenvalues in the diffusion operator. Following previous recommendations, we set *α* to 0.5 and t to 0 to maintain global relationships between data points in the embedded space. Setting t = 0 allows the diffusion time to be automatically estimated via a damped regularization process. Then, Procrustes rotation was used to align the obtained gradients to the average gradient of the typically-developed subjects exposed to the audiovisual stimulation. This procedure corrects for potential ambiguities in eigenvector ordering (such as sign flipping or changes in explained variance when eigenvalues are tied), ensuring that gradients computed separately for each individual are directly comparable and therefore improving the stability of the results. In order to assess whether the heterogeneity expressed along each gradient was altered we computed the numerical range of each of the first three gradient as the difference between the minimum and maximum eigenvector values. This metric reflects the degree of segregation, or in other words the distinct connectivity profiles, between the gradient extremes.

#### Network Based Statistic

To determine the statistically significant difference in FC between distinct groups we employed a networkbased statistical approach. This nonparametric statistical method controls family-wise error rates in network data by accounting for multiple comparisons. It first identifies connected components within the graph, based on edges that are significantly above the statistical threshold (F-contrast; here, an F-value threshold of 9, two-sided, with an alpha level of 0.05). The statistical significance of each connected component is then assessed by comparing its topology to a null distribution derived from nonparametric permutation testing of component sizes. By testing on a component-by-component basis, this approach achieves greater statistical power compared to mass-univariate methods. Our aim was to identify consistent changes across multiple comparisons excluding effects related to the temporary absence of a sensory modality. Following the approach by Luppi *et al*. [51], we employed a composite null hypothesis significance test, where the null hypothesis assumes that at least one of the datasets has no effect. Rejecting the null hypothesis requires that all comparisons show non-zero effects. This test compares an observed test statistic to a null distribution, with the test statistic defined as the minimum of the three F-scores from the relevant comparisons (e.g., audio video vs. blind, blind vs. audio, but not audio video vs. audio). To construct the null distribution, we randomly reshuffled (n=500) one dataset at a time and recalculated the F-scores. This selective reshuffling (of only one dataset at random rather than all) tests the observed data against the “least altered” version compatible with the null hypothesis. This method represents a least favorable configuration test, ensuring control over the false positive rate below a threshold of 0.05.

#### Brain correlation with film annotations

To establish how each ROI correlates with the audio, video, semantic, and speech movie features for each ROI in the cortical atlas, the Pearson correlation was computed between the corresponding BOLD signal and the features timeseries. When more than one feature was present for a given category, the maximum observed correlation was considered. With respect to the speech time series, to account for possible dependencies with the low-level auditory features, analyses were performed using partial Pearson correlation, regressing out the effects attributable to the low level sound properties, and thus retaining the correlation between the residuals and the BOLD signal. The statistical significance was corrected for False Discovery Rate (FDR) with BenjaminiHochberg (BH) correction [79] for all tests performed.

#### Brain maps correlation

The correlation between functional gradients was evaluated using Spearman’s rank-based, non-parametric correlation coefficient to ensure robustness against potential outliers. To mitigate potential biases from spatial autocorrelation and contralateral symmetry, we computed corrected p-values using an autocorrelation-preserving null model [56]. We simulated surrogate brain maps with spatial autocorrelation that matched the one of the original brain maps of interest that is functional gradients and movie parcellation. Using the geodetic distance matrix from the atlas as input, we generated 10,000 surrogate maps that preserve spatial autocorrelation. We then computed Spearman’s rank-based correlation for each of the obtained maps. The nonparametric p-values were computed considering the number of the correlation with surrogate maps that exceeded the observed correlation with the original data. For group comparisons, two-sample t-tests with Welch’s correction [80] were applied to account for unequal variances. Multiple testing corrections followed the FDR, with significance set at an alpha level of 0.05. Effect sizes were reported using Hedges’s g [81], a standardized measure that provides a less biased estimate of mean difference and that is especially useful for small sample sizes.

## Authors Contributions

## Acknowledgments

AIL acknowledges support from St John’s College, Cambridge; and a Wellcome Early Career Award (grant number 226924/Z/23/Z). GP acknowledges support from the European Research Council (ERC) Consolidator Grant under the European Union’s Horizon Europe programme (grant agreement No. 101171380, project RUNES).” MT is supported by PRIN 2022 (2022NEE53Z) from the Ministry of University and Research (MUR), the National Recovery and Resilience Plan – PNRR – “MNESYS” (PE00000006), with a specific contribution from the sub-project (“bando a cascata”) “SPARKS” (CUP D93C22000930002), and the ERC Proof of Concept “PRISM” (1011583). ER work was supported by the PRIN grants (20223K8B3X and P20228PHN2) by the Italian Ministry of University and Research and by the “Tuscany Health Ecosystem—THE” Project, Spoke 8, granted by Next Generation EU—National Recovery and Resilience Plan (Piano Nazionale di Ripresa e Resilienza, NRRP)—Mission 4 Component 2 Investment 1.4—Ministry of University and Research (MUR) Call N. 3277, Project Code ECS00000017 to E.R.

We would like to thank all the people behind the 101Dalmatians project. We thank Daniel Margulies, Marcin A. Radecki, Luca Cecchetti, Davide Bottari, Giacomo Handjaras and the colleagues for suggestions on the research project and data processing.

For the purpose of open access, the authors have applied a Creative Commons Attribution (CC BY) licence to any Author Accepted Manuscript version arising from this submission.

