## Supplemental information for "Beyond Reorganization: Intrinsic cortical hierarchies constrain experience-dependent plasticity in sensory-deprived humans"

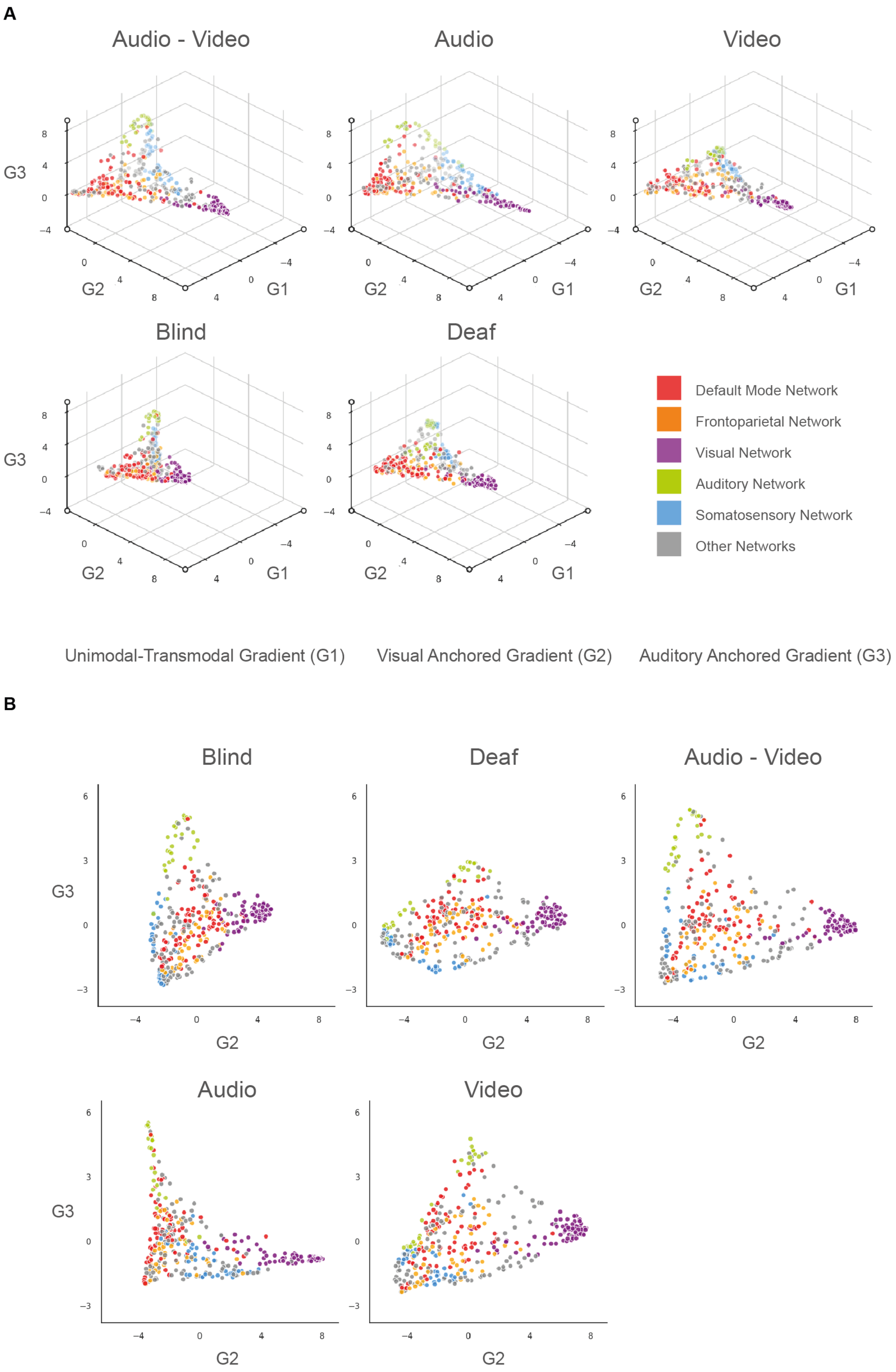


Figure S1. **Diffusion gradient space. A,** Three-dimensional representation of the first three gradients in typically developed individuals under audio-only, visual-only, and audiovisual stimulation. Each point represents a region from the multimodal HCP atlas [7], colored according to its primary functional network. **B,** Two-dimensional representation of the second and third gradients for the same conditions. Each point corresponds to a multimodal HCP atlas [7] region, with colors indicating the associated functional network.

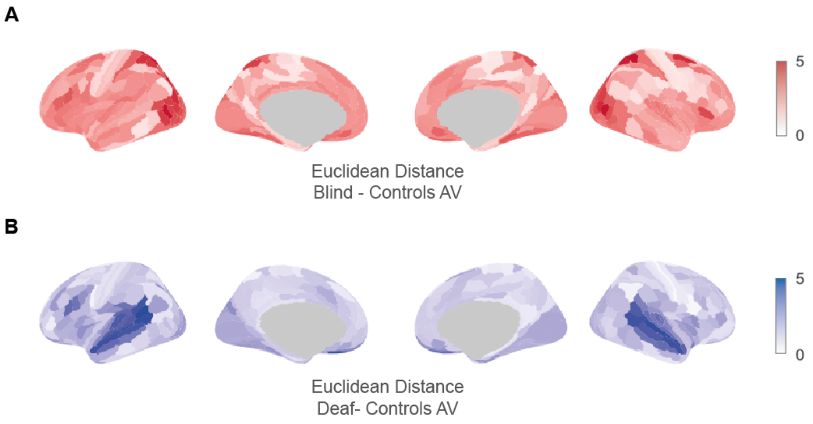


Figure S2. **Regional functional displacement. A,** Mean Euclidean distance in the three-dimensional diffusion embedding space (based on the first three gradients) for each region of the multimodal HCP atlas [7], comparing typically developed individuals under audiovisual stimulation with congenitally blind individuals. **B,** Same analysis as in A, comparing typically developed individuals with congenitally deaf individuals.

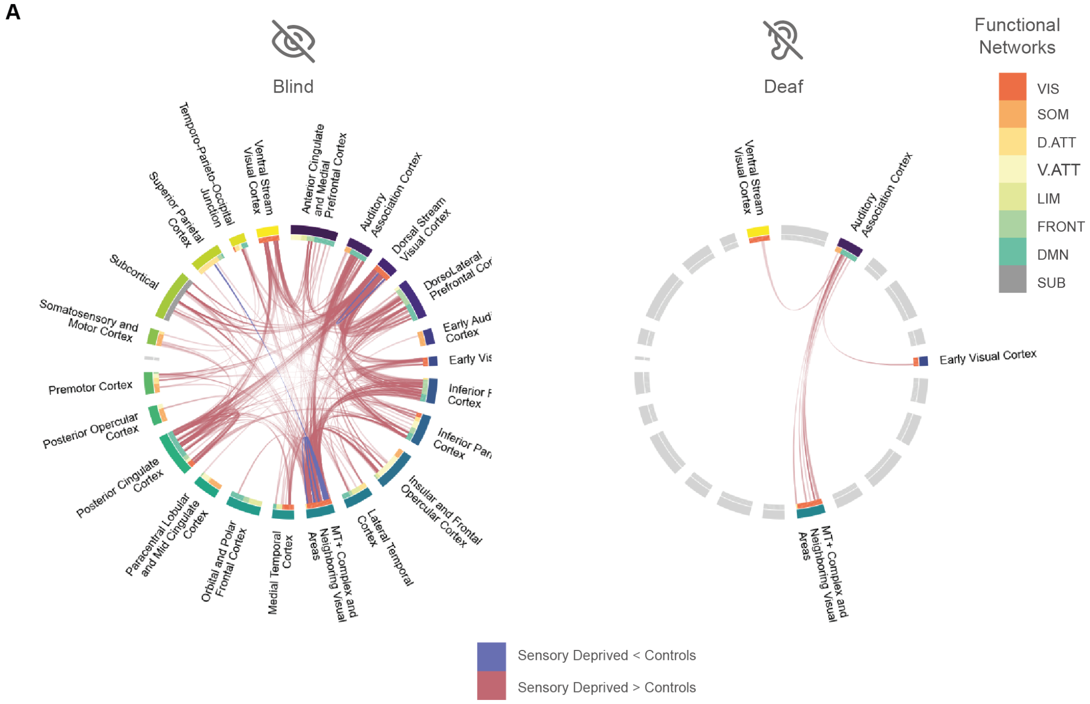


Figure S3. **Group-level differences in whole-brain functional connectivity. A,** Group differences in functional connectivity between congenitally blind or deaf individuals and matched typically developed controls identified using Network-Based Statistics (NBS; F $\geq$ 9, $p < 0.05$). Chord diagrams show significantly increased (blue) and decreased (red) connections for each sensory-deprived group. As in Figure 4, results are shown here according to the macro-anatomical subdivisions of the Glasser atlas. Congenitally blind individuals exhibited increased connectivity between deprived extrastriate visual areas (e.g., MT+ complex and dorsal stream cortices) and frontal regions (dorsolateral and inferior frontal cortex, spanning frontoparietal and DMN networks), alongside decreased intra-visual connectivity. Congenitally deaf individuals showed increased connectivity between visual areas (including MT+ and adjacent regions) and auditory association cortices (e.g., STS, A4, and A5).

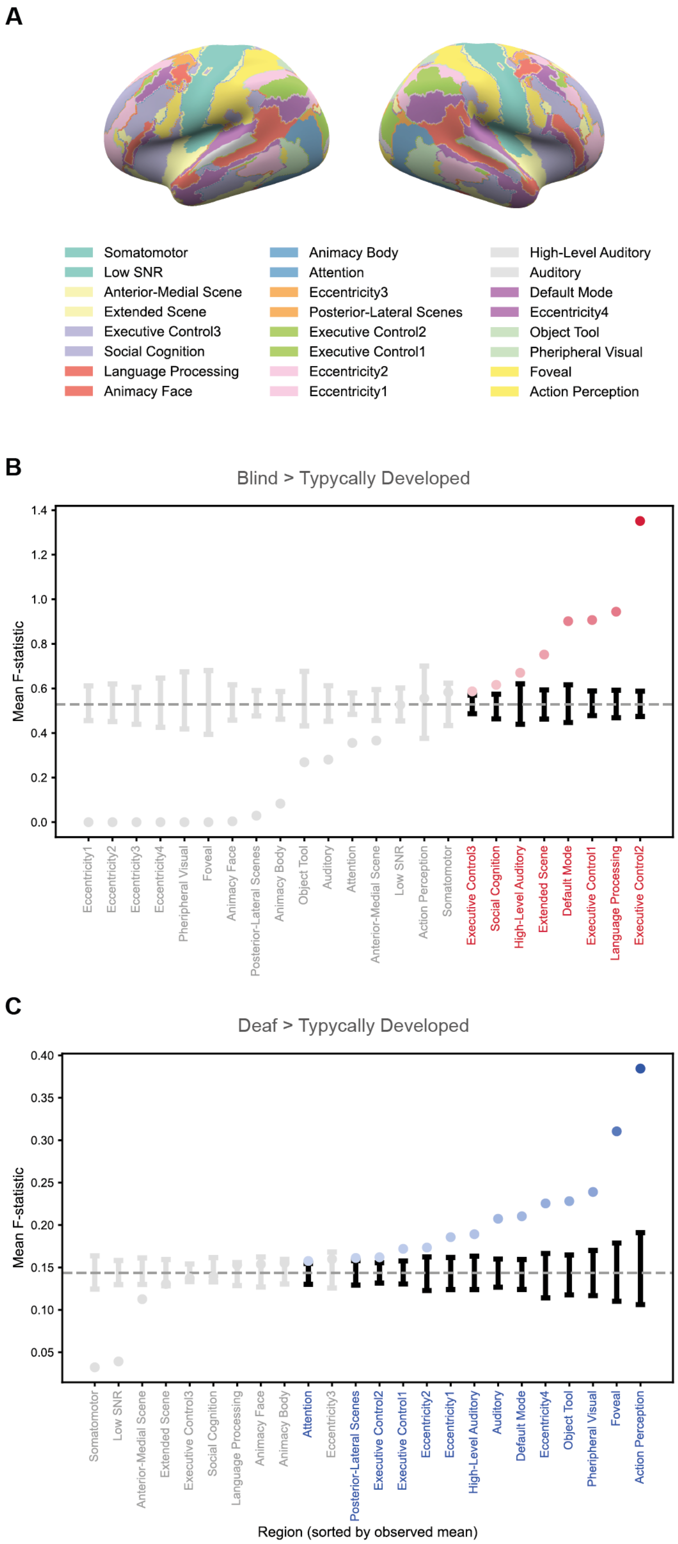


Figure S4. **Regional responses during film-watching across sensory-deprived groups. A,** Inflated cortical surface maps colored according to the functional parcellation defined by Rajimehr *et al.* [82], used to subdivide the cortex into parcels with consistent functional profiles during film viewing. **B, C,** To further characterize group differences observed during the film-watching task, we considered the mean composite F-statistic (as Figure 4) for each cortical region defined by Rajimehr *et al.* [82]. Functional terms are sorted by their observed mean F-statistic. Colored labels highlight regions where observed values exceed the 95th percentile of a null distribution generated from 10,000 spatial-autocorrelation-preserving surrogate maps [56], indicating statistically significant group effects. Grey points fall within the null distribution. Error bars represent the 95% confidence interval of the null distribution for each parcel.

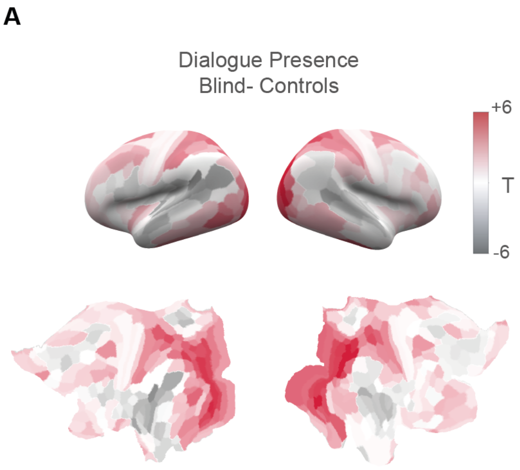


Figure S5. **Feature-specific recruitment of deprived sensory cortices. A,** BOLD time series from each cortical region were correlated with binary time courses of the presence or absence of dialogue. Correlation maps were generated per subject and compared across groups. Surface renderings and inflated ROI maps show results from region-wise two-tailed t-tests contrasting sensory-deprived and typically developed individuals. Blind participants show significantly stronger correlations between speech- related features and visual cortices (red, $p < 0.05$ FDR BH-corrected).

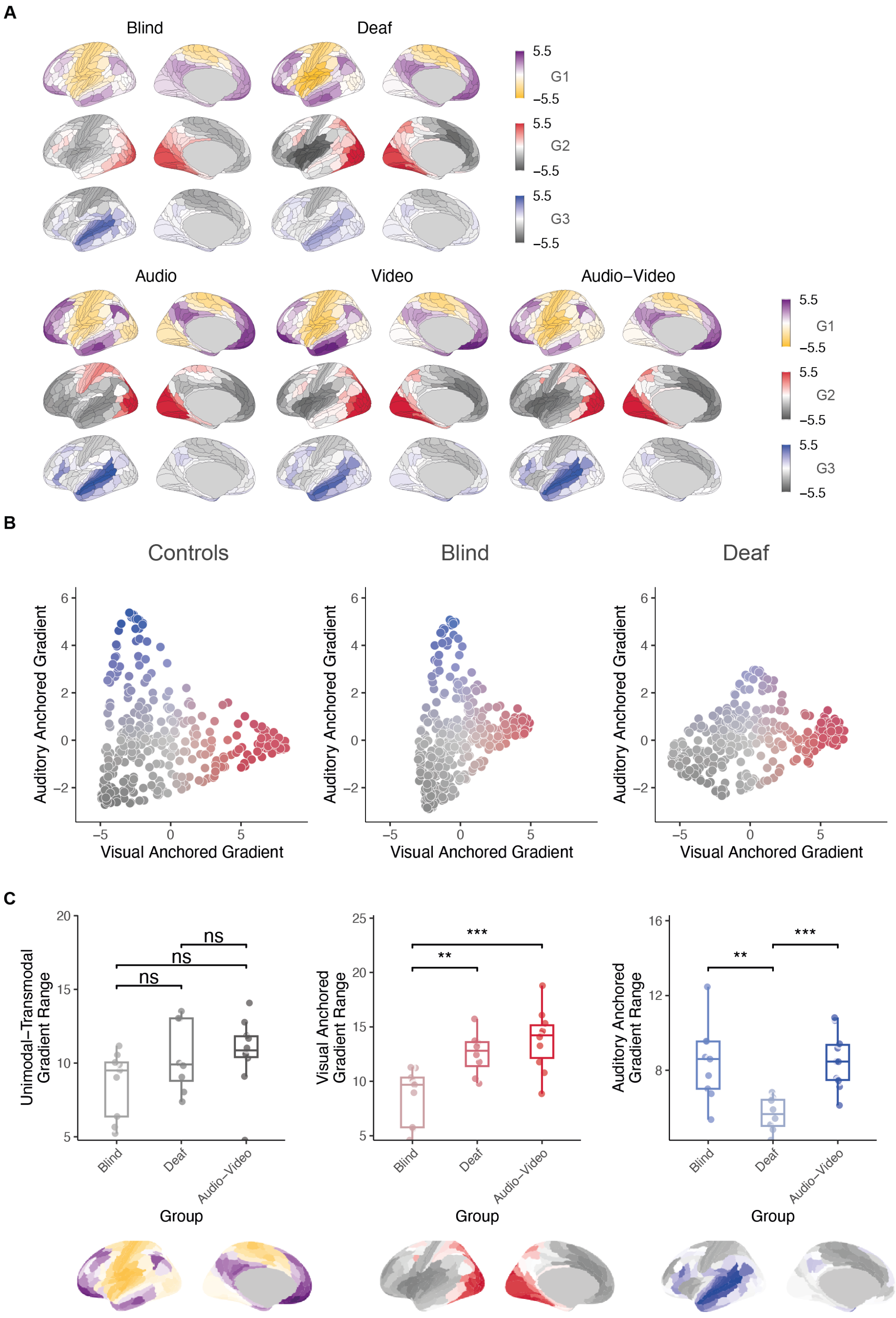


Figure S6. **Topographical maps and functional geometry under sensory deprivation (diffusion parameter $\alpha = 0.3$). A,** Surface maps of the first three cortical gradients in typically developed individuals under audio-only, visual-only, and audiovisual stimulation. Gradient 1 (G1) reflects the principal unimodal-to-transmodal axis, while G2 and G3 are anchored in visual and auditory cortices, respectively. Corresponding gradients in congenitally blind and deaf individuals, averaged across conditions. G1 remains consistent, whereas G2 and G3 show altered spatial structure, reflecting the lack of modality-specific input. **B,** Two-dimensional gradient embeddings illustrate compression along G2 in blind (horizontal arrow) and along G3 in deaf individuals (vertical arrow), relative to the audiovisual condition in controls. **C,** Boxplots show gradient range across groups. G1 does not differ significantly. Blind individuals show reduced G2 range; deaf individuals show reduced G3 range. Statistical tests are two-tailed t-tests (n.s. = not significant; ** $p < 0.01$; ***$p < 0.001$). Gradient topographies are shown below each plot.

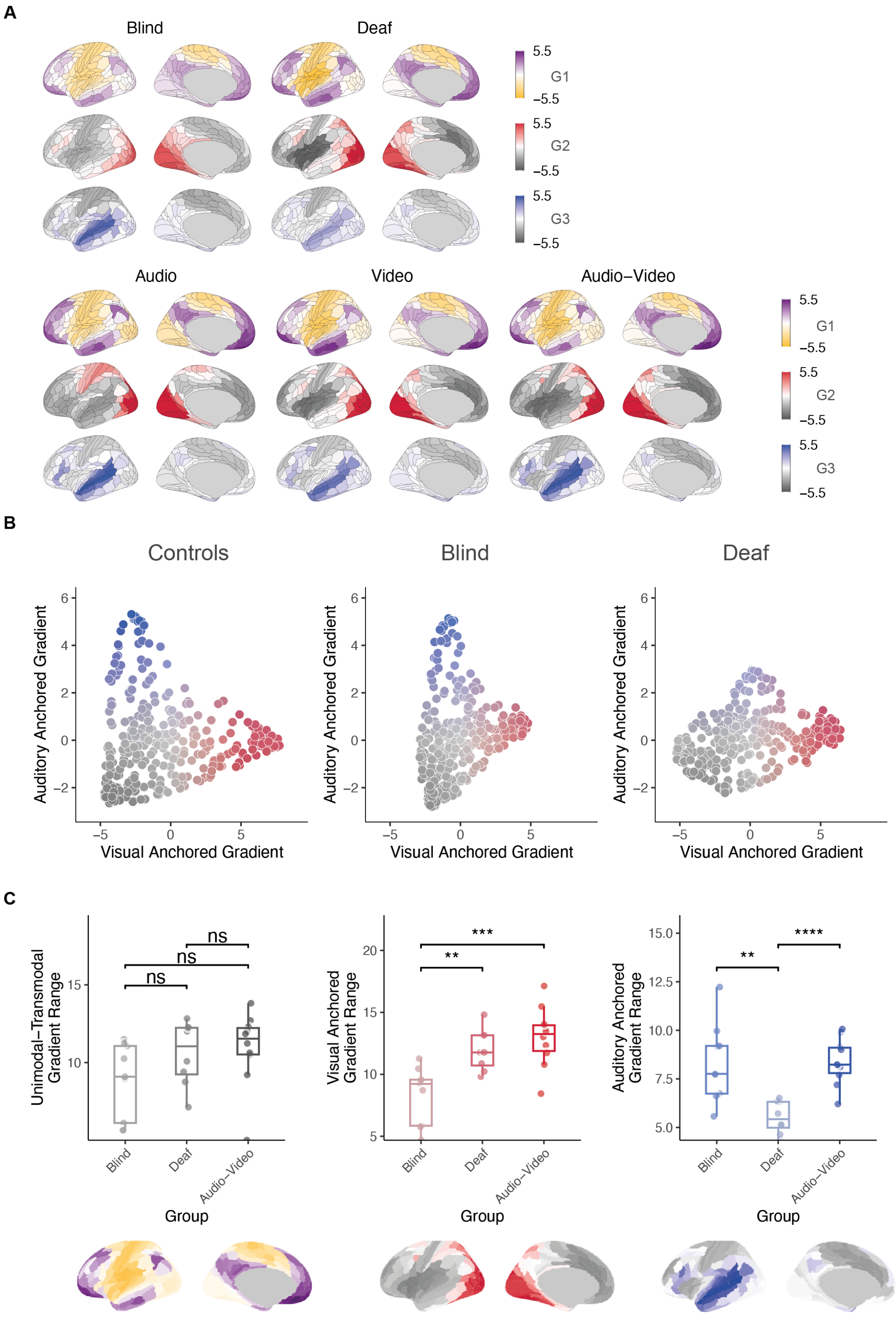


Figure S7. **Topographical maps and functional geometry under sensory deprivation (diffusion parameter $\alpha = 0.7$).A,** Surface maps of the first three cortical gradients in typically developed individuals under audio-only, visual-only, and audiovisual stimulation. Gradient 1 (G1) reflects the principal unimodal-to-transmodal axis, while G2 and G3 are anchored in visual and auditory cortices, respectively. Corresponding gradients in congenitally blind and deaf individuals, averaged across conditions. G1 remains consistent, whereas G2 and G3 show altered spatial structure, reflecting the lack of modality-specific input. **B,** Two-dimensional gradient embeddings illustrate compression along G2 in blind (horizontal arrow) and along G3 in deaf individuals (vertical arrow), relative to the audiovisual condition in controls. **C,** Boxplots show gradient range across groups. G1 does not differ significantly. Blind individuals show reduced G2 range; deaf individuals show reduced G3 range. Statistical tests are two-tailed t-tests (n.s. = not significant; ** $p < 0.01$; ***$p < 0.001$; ****$p < 0.0001$). Gradient topographies are shown below each plot.

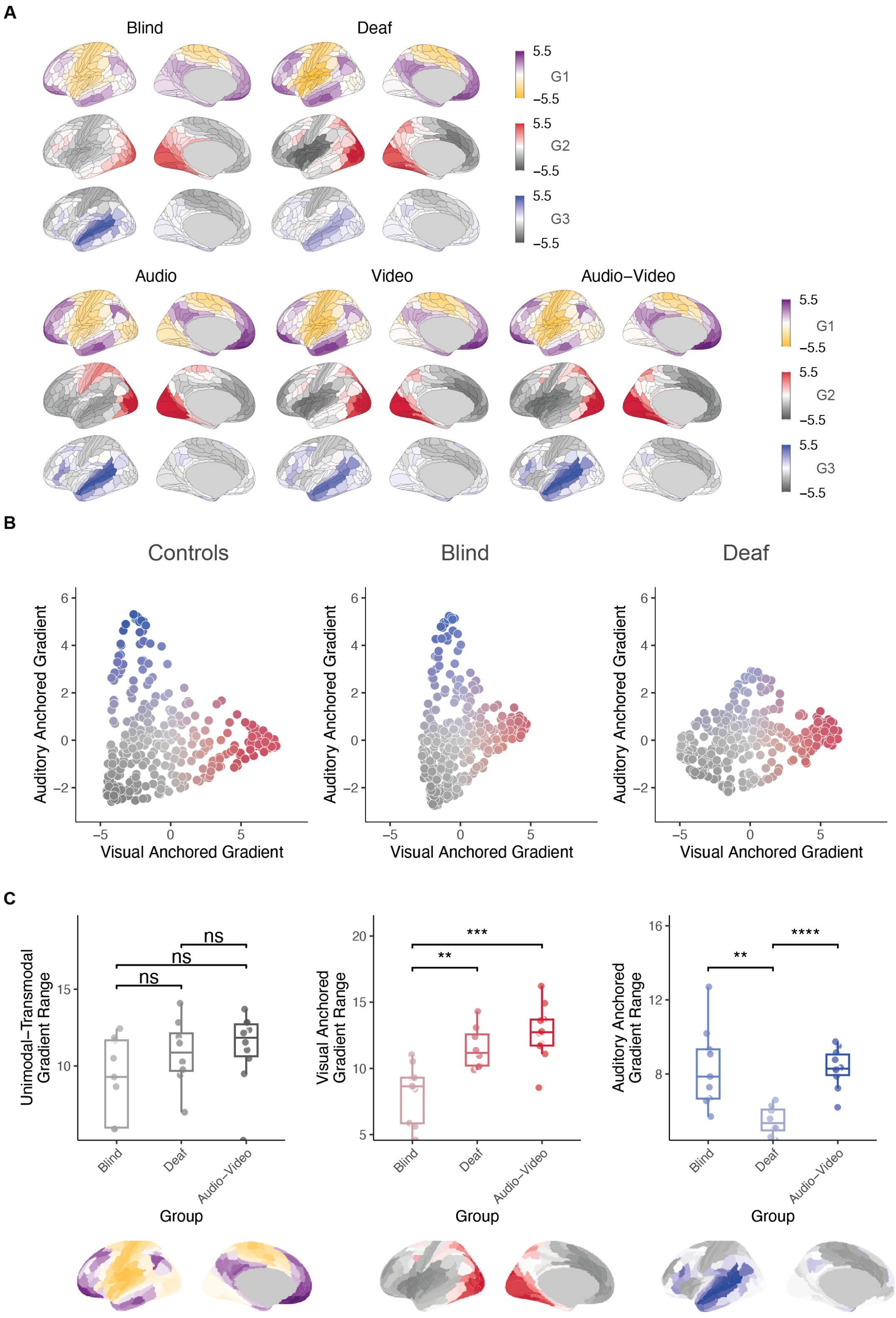


Figure S8. **Topographical maps and functional geometry under sensory deprivation (diffusion parameter $\alpha = 0.9$).A,** Surface maps of the first three cortical gradients in typically developed individuals under audio-only, visual-only, and audiovisual stimulation. Gradient 1 (G1) reflects the principal unimodal-to-transmodal axis, while G2 and G3 are anchored in visual and auditory cortices, respectively. Corresponding gradients in congenitally blind and deaf individuals, averaged across conditions. G1 remains consistent, whereas G2 and G3 show altered spatial structure, reflecting the lack of modality-specific input. **B**, Two-dimensional gradient embeddings illustrate compression along G2 in blind (horizontal arrow) and along G3 in deaf individuals (vertical arrow), relative to the audiovisual condition in controls. **C,** Boxplots show gradient range across groups. G1 does not differ significantly. Blind individuals show reduced G2 range; deaf individuals show reduced G3 range. Statistical tests are two-tailed t-tests (n.s. = not significant; ** $p < 0.01$; ***$p < 0.001$; ****$p < 0.0001$). Gradient topographies are shown below each plot.

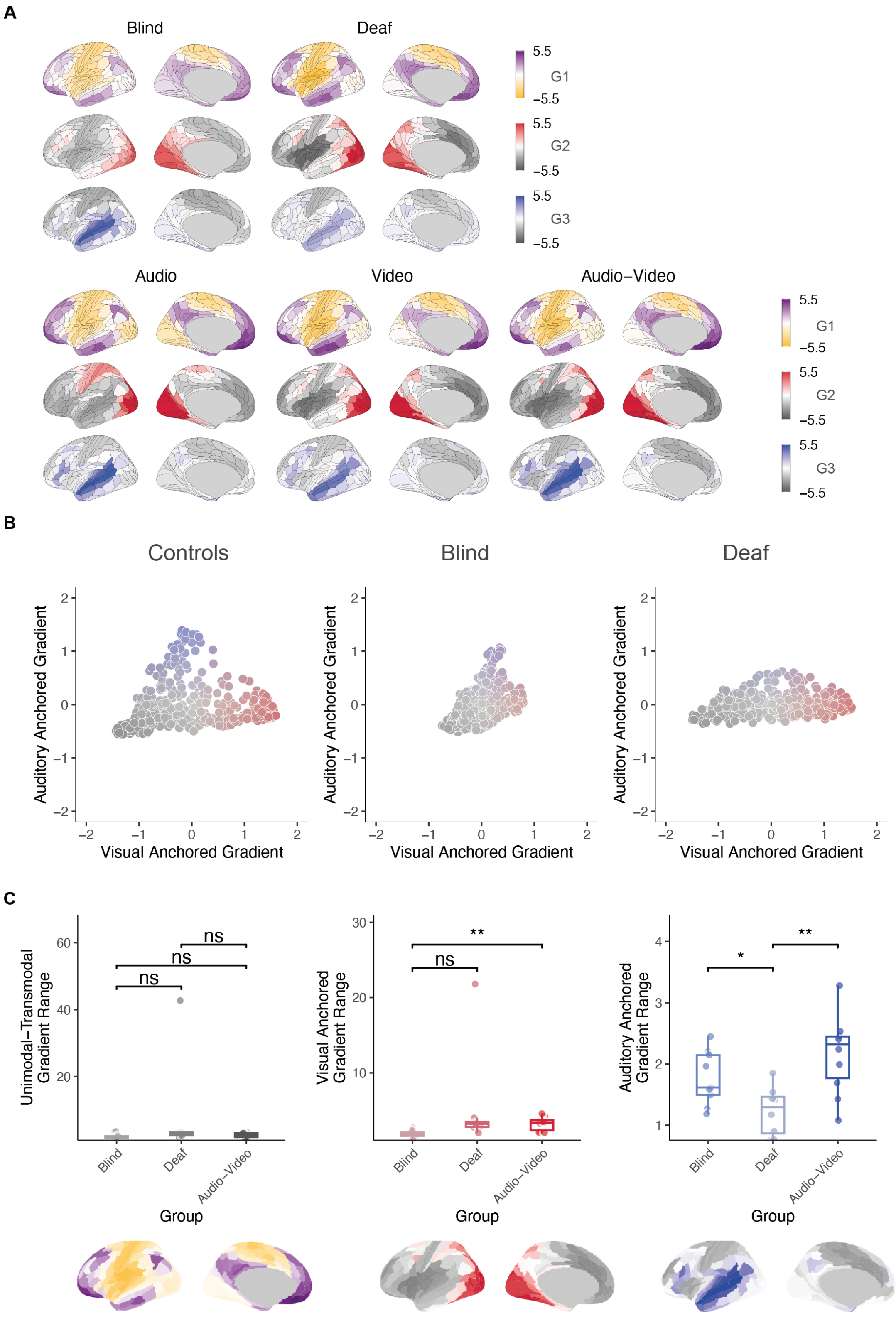


Figure S9. **Topographical maps and functional geometry under sensory deprivation (sparsity $\delta = 70\%$). A,** Surface maps of the first three cortical gradients in typically developed individuals under audio-only, visual-only, and audiovisual stimulation. Gradient 1 (G1) reflects the principal unimodal-to-transmodal axis, while G2 and G3 are anchored in visual and auditory cortices, respectively. Corresponding gradients in congenitally blind and deaf individuals, averaged across conditions. G1 remains consistent, whereas G2 and G3 show altered spatial structure, reflecting the lack of modality-specific input. **B,** Two-dimensional gradient embeddings illustrate compression along G2 in blind (horizontal arrow) and along G3 in deaf individuals (vertical arrow), relative to the audiovisual condition in controls. **C,** Boxplots show gradient range across groups. G1 does not differ significantly. Blind individuals show reduced G2 range; deaf individuals show reduced G3 range. Statistical tests are two-tailed t-tests (n.s. = not significant; $^*p < 0.05$; $^{**}\ p < 0.01$). Gradient topographies are shown below each plot.

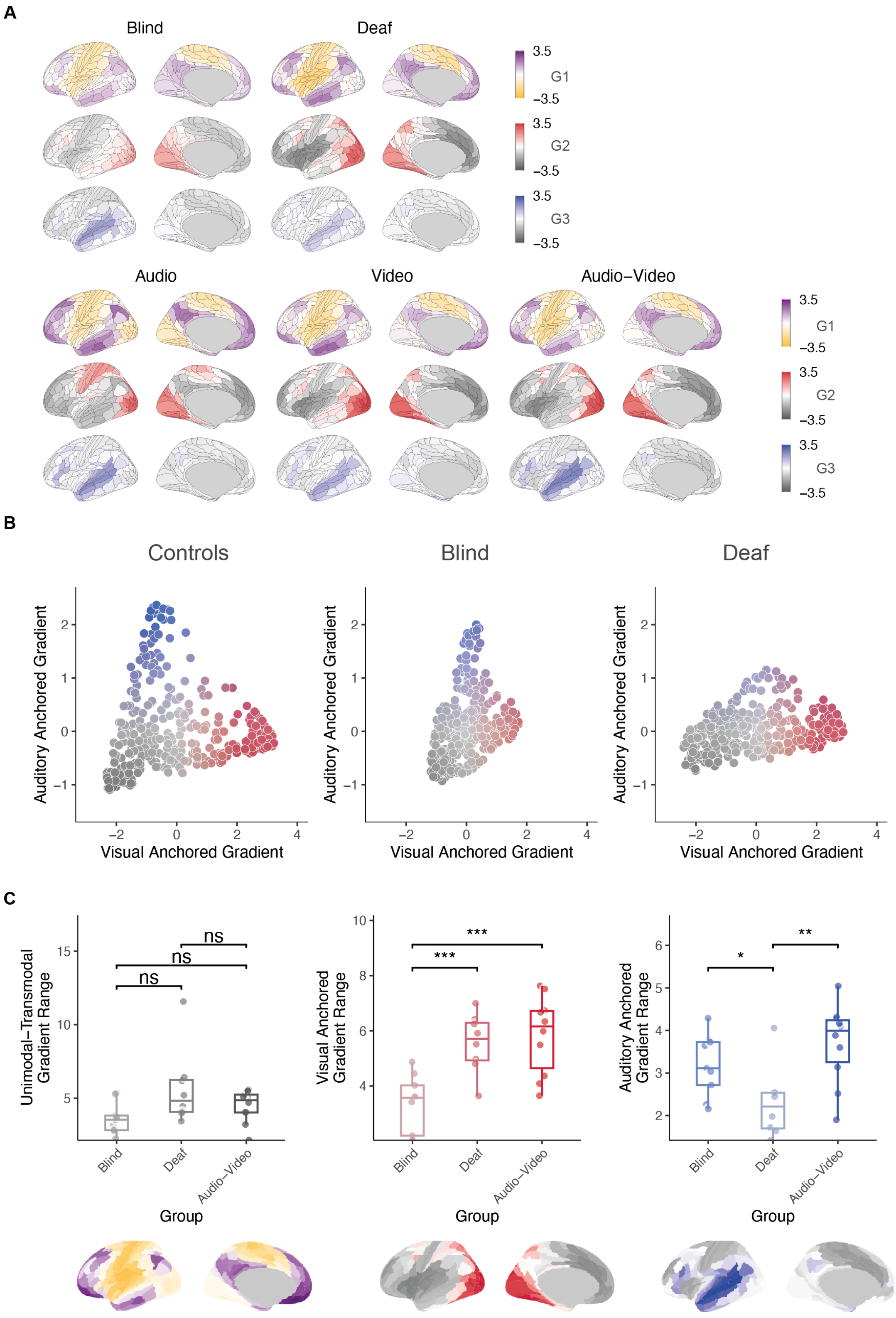


Figure S10. **Topographical maps and functional geometry under sensory deprivation (sparsity $\delta = 80\%$).A,** Surface maps of the first three cortical gradients in typically developed individuals under audio-only, visual-only, and audiovisual stimulation. Gradient 1 (G1) reflects the principal unimodal-to-transmodal axis, while G2 and G3 are anchored in visual and auditory cortices, respectively. Corresponding gradients in congenitally blind and deaf individuals, averaged across conditions. G1 remains consistent, whereas G2 and G3 show altered spatial structure, reflecting the lack of modality-specific input. **B,** Two-dimensional gradient embeddings illustrate compression along G2 in blind (horizontal arrow) and along G3 in deaf individuals (vertical arrow), relative to the audiovisual condition in controls. **C,** Boxplots show gradient range across groups. G1 does not differ significantly. Blind individuals show reduced G2 range; deaf individuals show reduced G3 range. Statistical tests are two-tailed t-tests (n.s. = not significant; $*p < 0.05$; $**\ p < 0.01$; $***\ p < 0.001$). Gradient topographies are shown below each plot.

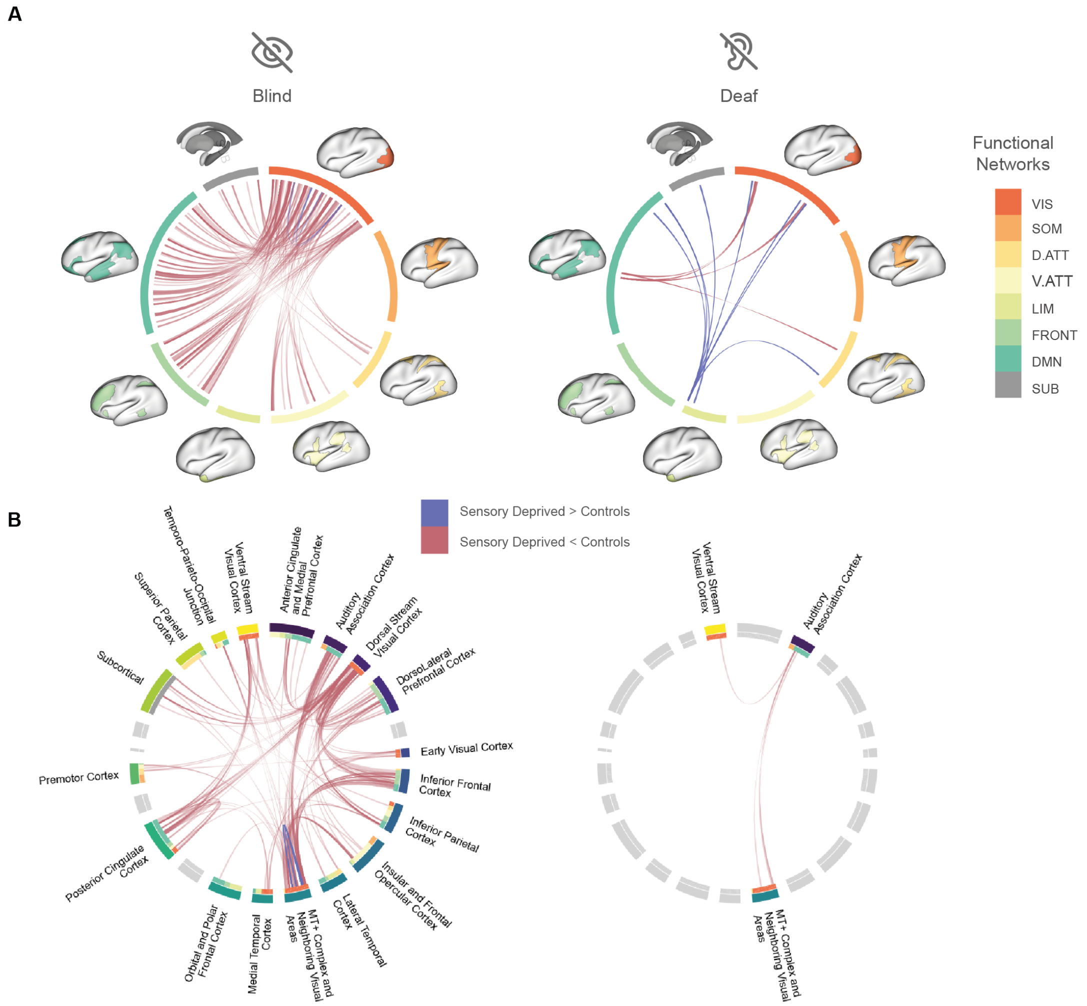


Figure S11. **Group-level differences in whole-brain functional connectivity (F¿12). A,** Group differences in functional connectivity between congenitally blind or deaf individuals and matched typically developed controls, identified using Network-Based Statistics with a more stringent threshold (NBS; F $\geq$ 12, $p < 0.05$). Chord diagrams display significantly increased (blue) and decreased (red) connections in each sensory-deprived group. Cortical nodes are grouped according to canonical functional networks [54]: VIS = visual, SOM = somatomotor, D.ATT = dorsal attention, V.ATT = ventral attention, LIM = limbic, FRONT = frontoparietal, DMN = default mode, SUB = subcortical. **B,** The same results are shown using macro-anatomical subdivisions from the Glasser atlas. Congenitally blind individuals exhibit increased connectivity between deprived extrastriate visual areas (e.g., MT+ complex and dorsal stream cortices) and frontal regions (dorsolateral and inferior frontal cortex, spanning frontoparietal and DMN networks), along with decreased intra-visual connectivity. Congenitally deaf individuals show increased connectivity between visual areas (including MT+ and adjacent regions) and auditory association cortices.

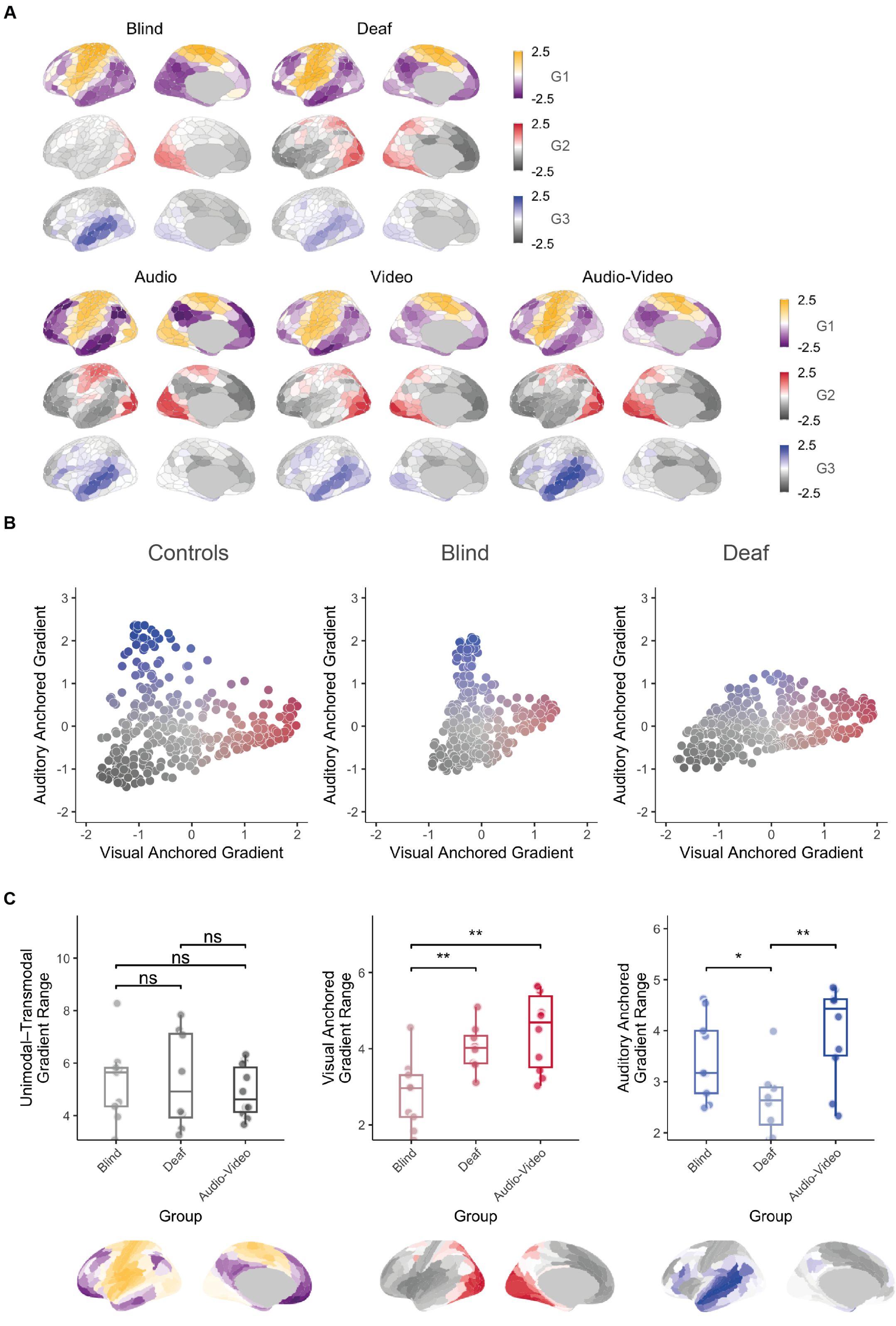


Figure S12. **Topographical maps and functional geometry under sensory deprivation (alternative parcellation Schaefer400). A,** Surface maps of the first three cortical gradients in typically developed individuals under audio-only, visual-only, and audiovisual stimulation. Gradient 1 (G1) reflects the principal unimodal-to-transmodal axis, while G2 and G3 are anchored in visual and auditory cortices, respectively. Corresponding gradients in congenitally blind and deaf individuals, averaged across conditions. G1 remains consistent, whereas G2 and G3 show altered spatial structure, reflecting the lack of modality-specific input. **B,** Two-dimensional gradient embeddings illustrate compression along G2 in blind (horizontal arrow) and along G3 in deaf individuals (vertical arrow), relative to the audiovisual condition in controls. **C,** Boxplots show gradient range across groups. G1 does not differ significantly. Blind individuals show reduced G2 range; deaf individuals show reduced G3 range. Statistical tests are two-tailed t-tests (n.s. = not significant; $*p < 0.05$; $**\ p < 0.01$). Gradient topographies are shown below each plot.
